# The proximal N-terminus of IRAG is required for potentiation of HCN4 channels

**DOI:** 10.64898/2026.04.20.719713

**Authors:** Lucas M. Blecker, Emily M. Teichman, Colin H. Peters, Daniel J. Enders, Rebecca Roth, William G. Nichols, Avery A. Langley, Cathy Proenza, John R. Bankston

**Author notes:** Corresponding Author John Bankston, PhD, Department of Physiology & Biophysics, University of Colorado – Anschutz Medical Campus, 12800 E. 19th Ave, MS 8307, Aurora, CO 80045.

## Abstract

The inositol triphosphate-associated, ER transmembrane proteins IRAG and LRMP are isoform specific regulators of the hyperpolarization-activated cyclic nucleotide-sensitive isoform 4 (HCN4) channel. LRMP prevents cAMP-dependent potentiation of HCN4, while IRAG mimics the effect of cAMP on the channel. We previously showed that regulation by LRMP requires both the N-terminus of HCN4 and a unique orientation of the HCN4 cAMP transduction center, which is comprised of the N-terminal HCN domain, the C-linker, and the S4-S5 linker. However, it remains unknown if the homologous IRAG requires similar structural features to mimic cAMP-dependent potentiation, or if the site and mechanism of action are different between the two regulators. Using patch clamp electrophysiology, we determined that the initial 43 amino acids of IRAG are necessary and sufficient to confer regulation of HCN4. Similar to LRMP, IRAG also requires a portion of the N-terminus of HCN4 to confer its regulatory effects. Also similar to LRMP, two point mutations in the C-linker region, which are the only sequence differences in that region between HCN4 and the other HCN isoforms, were able to eliminate the effect of IRAG suggesting the unique orientation of the cAMP transduction center in HCN4 is likely important for IRAG function. Taken together, these findings suggest a model whereby IRAG and LRMP interact with the channel in similar regions, although potentially in unique ways, and act on the cAMP transduction center with LRMP inhibiting the coupling of this region to gating and IRAG strengthening it.

**SUMMARY:** The ER transmembrane protein IRAG binds to and potentiates HCN4 channels. This study demonstrates that IRAG regulation of HCN4 requires only the first 43 amino acids of IRAG and involves contributions from the N-terminus and cAMP transduction center of HCN4.

## INTRODUCTION

The hyperpolarization-activated, cyclic nucleotide-sensitive (HCN) ion channels comprise a family of mixed cation channels that are activated at hyperpolarized potentials and potentiated by cyclic nucleotides (Santoro and Tibbs, 1999; DiFrancesco and Tortora, 1991). HCN channels are widely expressed throughout the body where they broadly act to maintain membrane excitability (Notomi and Shigemoto, 2004; Herrmann et al., 2015; Szegedi et al., 2023). HCN channels are homotetramers that share the subunit architecture of voltage gated K^+^ channels, with 6 transmembrane spanning domains (S1-S6) in each subunit and intracellular amino and carboxyl terminals (Lee and MacKinnon, 2017; Saponaro et al., 2021). There are four HCN isoforms (HCN1-4) that have unique gating kinetics and responses to cyclic nucleotides (Ludwig et al., 1998, 1999; Moosmang et al., 2001). The transmembrane domains, C-terminal cyclic nucleotide binding domain (CNBD), C-linker, and N-terminal HCN domain (HCND) of HCN channels are highly conserved; the major differences between isoforms reside in the distal portions of the N- and C-terminals.

Despite the structural similarities with voltage-gated K^+^ channels, HCN channels are activated by membrane hyperpolarization instead of depolarization. They are modulated by cyclic nucleotides via a conserved CNBD in the proximal C-terminus (Wang et al., 2001). Binding of cAMP potentiates HCN channels by shifting the voltage dependence of activation to more depolarized potentials, speeding the activation rate, and slowing the deactivation rate (Wicks et al., 2011; Wainger et al., 2001; Gross et al., 2018). While the domains required for the transduction of the cAMP binding to the channel gate are largely known, the detailed transduction mechanism is not completely understood. It is thought that the conversion of the signal that turns cAMP binding into a change in channel gating occurs through interactions within the ‘CNBD transduction center’ (Porro et al., 2019; Wang et al., 2020; Sunkara et al., 2018), comprised of the HCND, the S4-S5 intracellular loop, the CNBD, and the C-linker that connects S6 with the CNBD (Lee and MacKinnon, 2017; Saponaro et al., 2021).

We recently discovered a pair of homologous proteins that specifically modulate only the HCN4 isoform (Peters et al., 2020a, 2022). IRAG (Inositol triphosphate Receptor-Associated Guanylate kinase substrate; IRAG1; MRVI1) and LRMP (Lymphoid Restricted Membrane Protein; IRAG2, Jaw1) are homologous proteins with a shared overall architecture consisting broadly of four domains: a large cytoplasmic N-terminal region that includes a relatively well-conserved cytoplasmic coiled-coil domain, an SR/ER transmembrane segment, and a small C-terminal SR/ER luminal domain. The coiled-coil region is critical for the interactions of LRMP and IRAG with IP_3_Rs that regulate Ca^2+^ release from the ER in smooth muscle cells (Schlossmann et al., 2000; Geiselhöringer et al., 2004b; a; Prüschenk et al., 2021).

Despite their homology, LRMP and IRAG have opposing effects on HCN4 activation (Peters et al., 2020a). IRAG depolarizes the voltage dependence of activation of HCN4 in the absence of cAMP, while LRMP attenuates the ability of cAMP to shift the voltage dependence (Peters et al., 2020a). Interactions between HCN4 and IRAG or LRMP do not alter channel deactivation, nor do they prevent the slowing of channel deactivation that occurs upon cAMP binding, suggesting that neither prevents cAMP from binding to the CNBD (Peters et al., 2020a). Thus, their effects appear to result from aspects of the intramolecular cAMP signal transduction pathway that are unique to the HCN4 isoform.

We previously showed that regulation of HCN4 by LRMP requires the N-terminals of both LRMP and HCN4 as well as a unique orientation of the cAMP transduction center (Peters et al., 2024). However, it is not known if IRAG requires similar structural features to mimic cAMP-dependent potentiation, or if the sites and mechanisms of action are different between the two regulators. The major homology between the LRMP and IRAG lies within the coiled-coil regions and the ER transmembrane segments. Hence, it is possible that the unique portions of IRAG act on HCN4 via a different mechanism than LRMP. Conversely, it is possible that these two proteins share variations of a common mechanism with reversed polarity, such that LRMP prevents the coupling between cAMP binding and gating of HCN4 while IRAG mimics that coupling. To define the domains involved in IRAG regulation of HCN4, we used patch clamp electrophysiology and systematically truncated or mutated the various intracellular domains of both HCN4 and IRAG. We show here that the initial 43 amino acids of IRAG and the first 25 amino acids of HCN4 are necessary for IRAG regulation of HCN4 channels. In addition, mutation of 2 residues in the C-linker of HCN4 eliminate regulation by IRAG, suggesting that the unique orientation of the cAMP transduction center is critical for the IRAG effect, as it is for regulation by LRMP. These results are consistent with a model in which LRMP and IRAG extend up from the ER towards the plasma membrane, allowing their N-terminals to interact with the N-terminus of HCN4 to alter the function and polarity of the cAMP transduction center. These results are consistent with a model in which the opposing effects of LRMP and IRAG on HCN4 arises via subtle differences in polarity within a common overall mechanism.

## MATERIALS AND METHODS

### Cell culture, transfection, and peptides

HEK293 cells were obtained from ATCC (Manassas, VA), which uses STR profiling for cell line authentication. HEK 293 cells from which new cell lines were established and HEK HCN4 cells were negative for mycoplasma infection. Testing for mycoplasma infection was performed at the Molecular Biology Core Facility in the Barbara Center for Childhood Diabetes at the University of Colorado Anschutz Medical Campus. None of the cells are on the list of commonly misidentified cell lines.

HEK 293 cells were grown in a humidified incubator at 37 °C and 5% CO_2_ in high glucose DMEM with L-glutamine supplemented with 10% FBS, 100 U/mL penicillin, and 100 μg/mL streptomycin. Cells were transfected 48 hours prior to experiments and were plated on protamine-coated glass coverslips.

Patch clamp experiments were performed in either transiently transfected HEK293 cells, an HCN4 stable line in HEK293 cells (Zong et al., 2012), or a stable line in HEK293 cells of double mutant HCN4 P545A/T547F (HCN4 PT/AF). Stable cell lines were made by transfecting HEK293 cells with the respective plasmids using Lipofectamine 2000 (Invitrogen, Waltham, MA) according to the manufacturer’s instructions. Forty-eight hours post-transfection, 200 μg/mL of G418 disulfate (Alfa Aesar, Haverhill, MA) or Hygromycin B (InvivoGen, San Diego, CA) was added to the cell culture media in place of pen-strep to select for stably transfected cells. Single-cell clones were evaluated using whole-cell patch clamp and the clonal lines that exhibited the largest and most consistent currents were grown into stable cell lines.

Transient transfection of HCN4 constructs and/or IRAG was performed using Fugene6 (Promega, Madison, WI) or JetPrime (Polyplus-Sartorius, Illkirch, France), both using manufacturer’s instructions. Transfections of all constructs that did not include fluorescent tags were performed with the addition of eGFP as a co-transfection marker. All data were collected from a minimum of 3 transfections per condition. The n-values listed in the supplementary tables represent the number of individual cells that were patch-clamped for a given condition.

IRAG 1-43 peptide was manufactured by Lifetein LLC (Somerset, NJ) at >95% purity and included C-terminal amidation. Peptides were delivered as lyophilized powder, reconstituted in patch clamp intracellular solution, aliquoted, and kept frozen at -80 C°. Samples were used within 1 year of manufacture to maintain peptide quality. The specific amino acid sequence was: MGRSLTCPFGISPACGAQASWSIFGVGTAEVPGTHSHSNQAAA.

### Patch clamp electrophysiology

Cells were plated on sterile ECL-coated glass coverslips 24–48 hours prior to experiments. Cells on coverslip shards were transferred to the recording chamber and perfused (∼0.5–1 mL/min) with extracellular solution containing (in mM): 30 KCl, 115 NaCl, 1 MgCl_2_, 1.8 CaCl_2_, 5.5 glucose, and 5 HEPES. Transiently transfected cells were identified by green fluorescence.

Patch pipettes were pulled from borosilicate glass to a resistance of 1.0–3.0 MOhm when filled with intracellular solution containing (in mM): 130 K-Aspartate, 10 NaCl, 1 EGTA, 0.5 MgCl_2_, 5 HEPES, and 2 Mg-ATP. One mM cAMP was added to the intracellular solution as indicated. All recordings were performed at room temperature in the whole-cell configuration. Data were acquired at 5 KHz, and low-pass filtered at 10 KHz using an Axopatch 200B amplifier (Molecular Devices, San Jose, CA), Digidata 1440 A A/D converter and Clampex software (Molecular Devices). Pipette capacitance was compensated in all recordings. Membrane capacitance and series resistance (Rs) were estimated in whole-cell experiments using 5 mV test pulses. Only cells with a stable Rs of <10 MOhm were analyzed and currents at -130 mV > 200 pA. For HCN4-S719X data only, due to abnormally large current amplitudes, a series resistance compensation was used at 80:20 ratio of prediction and correction. Data were analyzed in Clampfit 10.7 (Molecular Devices).

Channel activation was determined from peak tail current amplitudes at –50 mV following 3 s hyperpolarizing pulses to membrane potentials between –50 mV and –170 mV from a holding potential of 0 mV. Normalized tail current-voltage relationships were fit by single Boltzmann equations to yield values for the midpoint activation voltage (V_1/2_) and slope factor. All reported voltages are corrected for a calculated +14 mV liquid junction potential between the extracellular and intracellular solutions.

### Statistical Analysis

All statistical analysis and graphing was performed using SigmaPlot 12.0 (Grafiti LLC, Palo Alto, CA), GraphPad Prism (GraphPad Software, Boston, MA), LabPlot (KDE, open source), and Adobe Illustrator (Adobe, San Jose, CA). Normality was assessed using the Shapiro-Wilk test. To prevent biasing of the results, all data were included except for cells showing large changes in leak or access resistance during the recording, those for which the access resistance was >10 MΩ at any point during recording, or those with currents less than 200 pA at -130 mV. Tests for differences in the average midpoint of activation for a given HCN channel construct in the presence of IRAG and/or 1 mM cAMP were performed with a 2-way ANOVA. A Bonferroni post-hoc test was used to test for multiple comparisons. The main independent variables were the absence or presence of IRAG and the absence or presence of 1 mM cAMP in the pipette solution. Only for HCN4 V604X, a student t-test was performed to test the differences in the average midpoint of activation in the presence or absence of IRAG 1-43-cit. All p values are reported in supplementary data tables.

## RESULTS

### The N-terminus of IRAG is required for regulation of HCN4

We previously demonstrated that co-expression of IRAG with HCN4, but not HCN1 or HCN2, results in a depolarizing shift in the voltage dependence of activation in the absence of cAMP and no additional shift in the presence of cAMP. To confirm that IRAG modulates HCN4 as we previously described and to provide a contemporaneous control dataset for further studies, we performed whole cell patch clamp electrophysiology using HEK 293 cells stably transfected with mouse HCN4 in the presence or absence of transiently transfected IRAG. We elicited currents by stepping from a holding potential of 0 mV down to successively more hyperpolarized potentials in 3 s voltage steps, and then stepping to -50 mV to elicit deactivating tail currents (**Fig. 1A**). We plotted peak tail current amplitudes as a function of activating voltage and fit the data with the Boltzmann equation to create conductance-voltage (GV) curves, from which we determined the V½ (**Fig. 1B)**. To determine the channel response to cyclic nucleotides, we measured HCN4 channels with 1 mM cAMP included in the patch pipette. As expected, inclusion of cAMP resulted in a ∼12 mV depolarizing shift in the voltage dependence of activation. To determine the effect of IRAG, we repeated each of these measures with full-length IRAG transiently expressed in our HEK cells that stably express HCN4. Consistent with our previous report (Peters et al., 2020a), we found that IRAG shifted the voltage dependence of HCN4 activation by ∼12 mV to more depolarized potentials and that cAMP did not produce an additional effect in the presence of IRAG (**Fig. 1B, C**).

**Figure 1.**
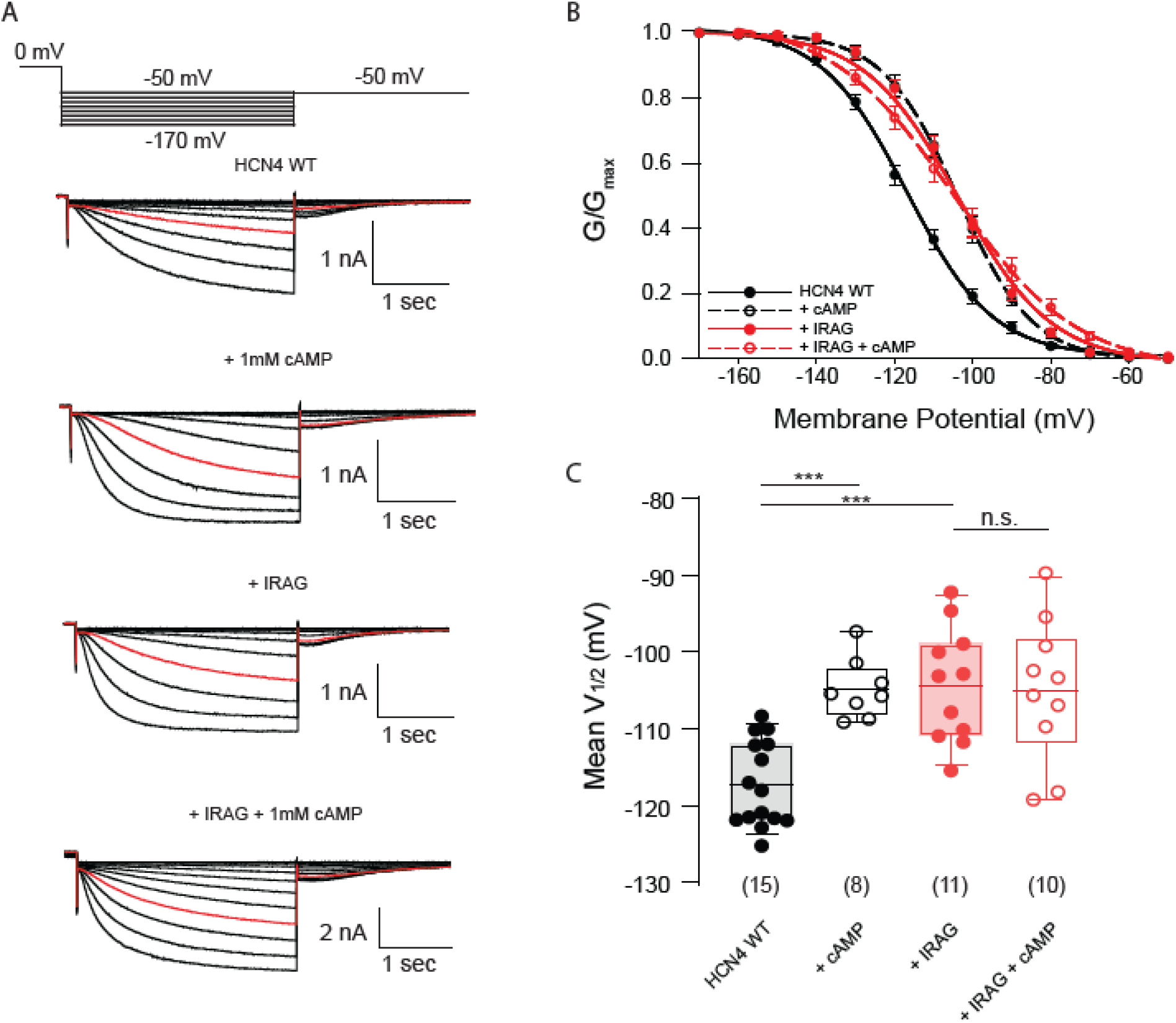
Full length IRAG potentiates HCN4. (A) Voltage clamp protocol used in all experiments in the present study (*top*). Currents from HEK293 cells stably expressing HCN4 in the presence or absence of 1 mM cAMP as well as in the presence or absence of IRAG, or both (*bottom*). Currents in red are elicited at -110 mV. (B) Voltage dependence of activation shown as normalized conductance vs. voltage (GV) curves for each condition averaged across cells; HCN4 alone (black) with and without 1 mM cAMP (open vs. colored circles) and with and without IRAG (red). Error bars represent ± standard error of the mean. (C) Box plot of mean midpoints of activation for HCN4 +/-IRAG and/or 1mM cAMP using the same color scheme as B. Lines represent mean, box represents IQR, whiskers represent 10-90 percentile range. * represents p < 0.05, ** represents p < 0.01, and *** represents p < 0.001. All means, standard errors, and exact p-values are in **Supplementary Table 1**.

We next sought to determine which domains of IRAG and HCN4 are required for regulation. IRAG is an 899 amino acid protein with a short C-terminal ER transmembrane domain and a large cytoplasmic N-terminal region with a coiled-coil domain that shares 60% similarity and 43% identity with LRMP in that region (**Fig. 2A, IRAG sequence: uniprot Q9WUX5, LRMP sequence: uniport:Q60664**). Our plan was to serially truncate IRAG starting from the N-terminal region of the protein by cutting ∼50 amino acids at a time in order to identify the regions that were critical. To our surprise, the first truncated IRAG construct we generated, which lacks the first 43 amino acids (IRAG Δ1-43), no longer shifted the voltage dependence of activation of HCN4 (**Fig. 2B, E**). Co-expression of IRAG Δ1-43 with HCN4 resulted in no shift in the V_1/2_. Addition of cAMP shifted channel activation to a similar degree in the presence or absence of IRAG Δ1-43. This lack of regulation could be because the first 43 amino acids are critical for the regulation. However, it is also possible that the truncation causes the protein to misfold, or that the protein fails to express. To determine whether residues 1-43 of IRAG are critical for regulation of HCN4, we expressed them as a plasmid with a C-terminal mCitrine fused to the peptide (IRAG 1-43-cit) to allow us to visualize expressing cells. Remarkably, IRAG 1-43-cit fully recapitulated the IRAG effect – it caused a significant depolarization in the V_1/2_ of HCN4, and cAMP had no significant effect on the V_1/2_ in the presence of IRAG 1-43-cit (**Fig. 2C, E**).

**Figure 2.**
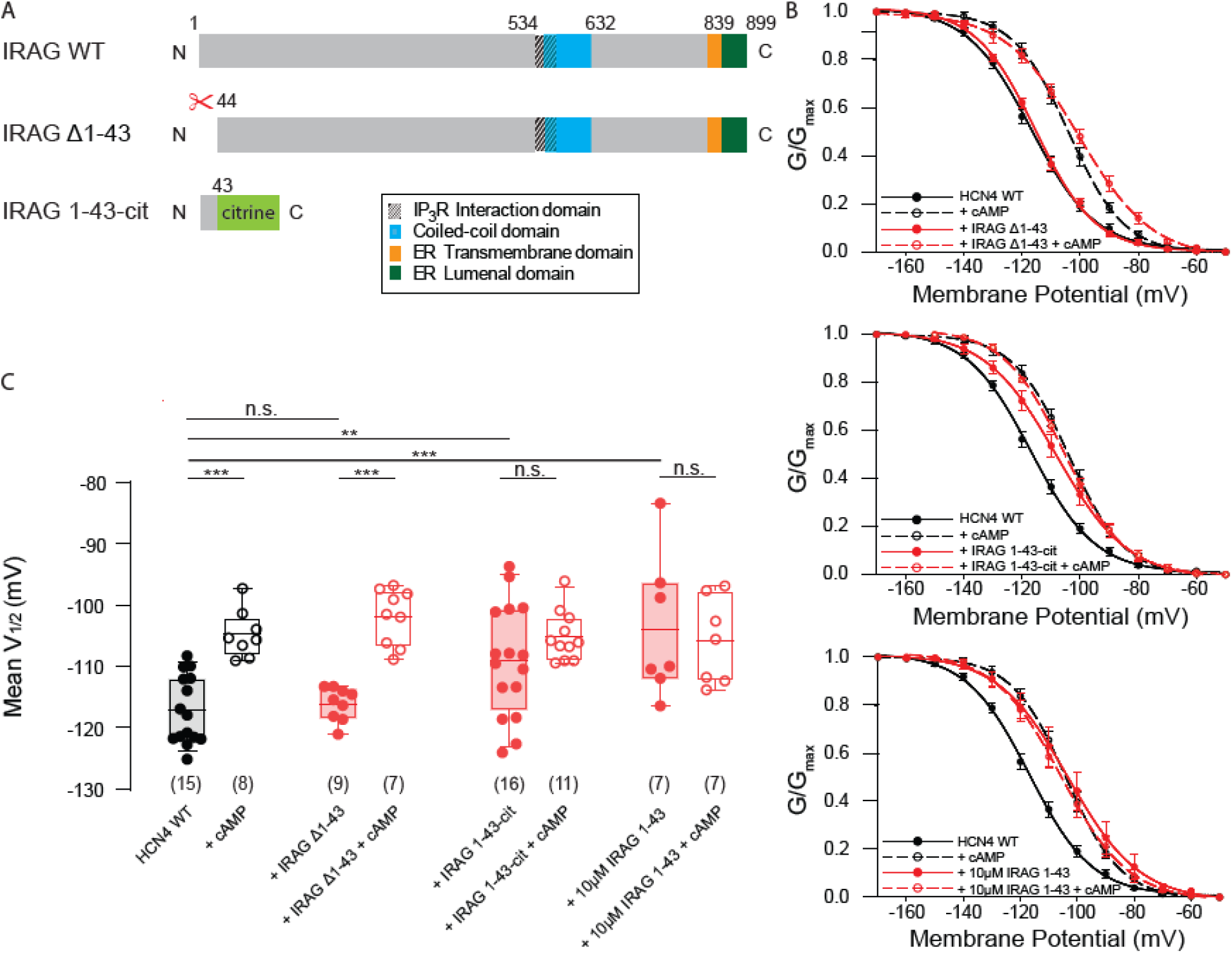
IRAG truncations show IRAG residues 1-43 are necessary and sufficient for IRAG regulation of HCN4. **(A)** Schematic depicting IRAG and its known structural domains, compared to two fragments tested in this figure for regulation of HCN4. **(B)** Voltage dependence of activation curves for HEK293 cells stably expressing HCN4 in the absence (black) or presence of either IRAG Δ1-43, IRAG 1-43-cit, or 10 μM IRAG 1-43 peptide (all +IRAG conditions in red) and/or 1 mM cAMP (open symbols). Error bars represent SEM. **(C)** Box plot of mean midpoints of activation for HEK293 cells stably expressing HCN4 transiently transfected with IRAG Δ1-43, IRAG 1-43-cit, or 10 μM IRAG 1-43 peptide in the absence or presence of 1 mM cAMP using the same color scheme as B. Lines represent mean, box represents IQR, whiskers represent 10-90 percentile range. * represents p < 0.05, ** represents p < 0.01, and *** represents p < 0.001. All means, standard errors, and exact p-values are in **Supplementary Table 2**.

Given this short N-terminal sequence of IRAG is sufficient to confer the full effect of IRAG on HCN4, we sought to confirm this result in the absence of the large fluorescent protein tag and without overexpression. To do this, we synthesized a peptide of the first 43 amino acids of IRAG and included the peptide in our internal patch solutions at 10 μM. Once again, the presence of this IRAG peptide was sufficient to cause the full shift in the V_1/2_ and the inclusion of cAMP in the pipette did not result in any additional changes (**Fig. 2D, E**). Together, these data suggest that the first 43 amino acids of IRAG are both necessary and sufficient to potentiate HCN4.

### The unique distal N- and C-termini of HCN4 contribute to IRAG regulation of HCN4

Given that IRAG modulates only the HCN4 isoform, that the bulk of IRAG is in the cytosol, and that a cytosolic fragment is sufficient to regulate HCN4, it is likely that IRAG interacts with the unique intracellular domains of HCN4. While the C-linker, the CNBD, and the HCND of HCN4 are highly conserved among HCN channel isoforms, the distal N- and C-termini are unique (**Fig. 3A**). Thus, we began probing HCN4 regions that may confer sensitivity to IRAG by examining the distal N- and C-termini. Previous work has shown that a naturally occurring alternative translation initiation site in the N-terminus of HCN4 removed the first 25 amino acids of the channel (Liu and Aldrich, 2011). For these experiments we transiently transfected our mutant channels with and without IRAG 1-43-cit and again measured currents in response to a series of voltage steps. HCN4Δ1-25 responded to 1 mM cAMP with a ∼9 mV depolarizing shift in voltage dependence (**Supplementary Table 3**). However, it failed to respond to co-expression with IRAG1-43-cit, showing no change in the voltage dependence of activation (**Fig. 3B, C**). Furthermore, addition of 1 mM cAMP to HCN4Δ1-25 channels in the presence of IRAG 1-43-cit resulted in a typical depolarizing shift in channel opening. This suggests that the distal N-terminus of HCN4 is required for IRAG potentiation of the channel.

**Figure 3.**
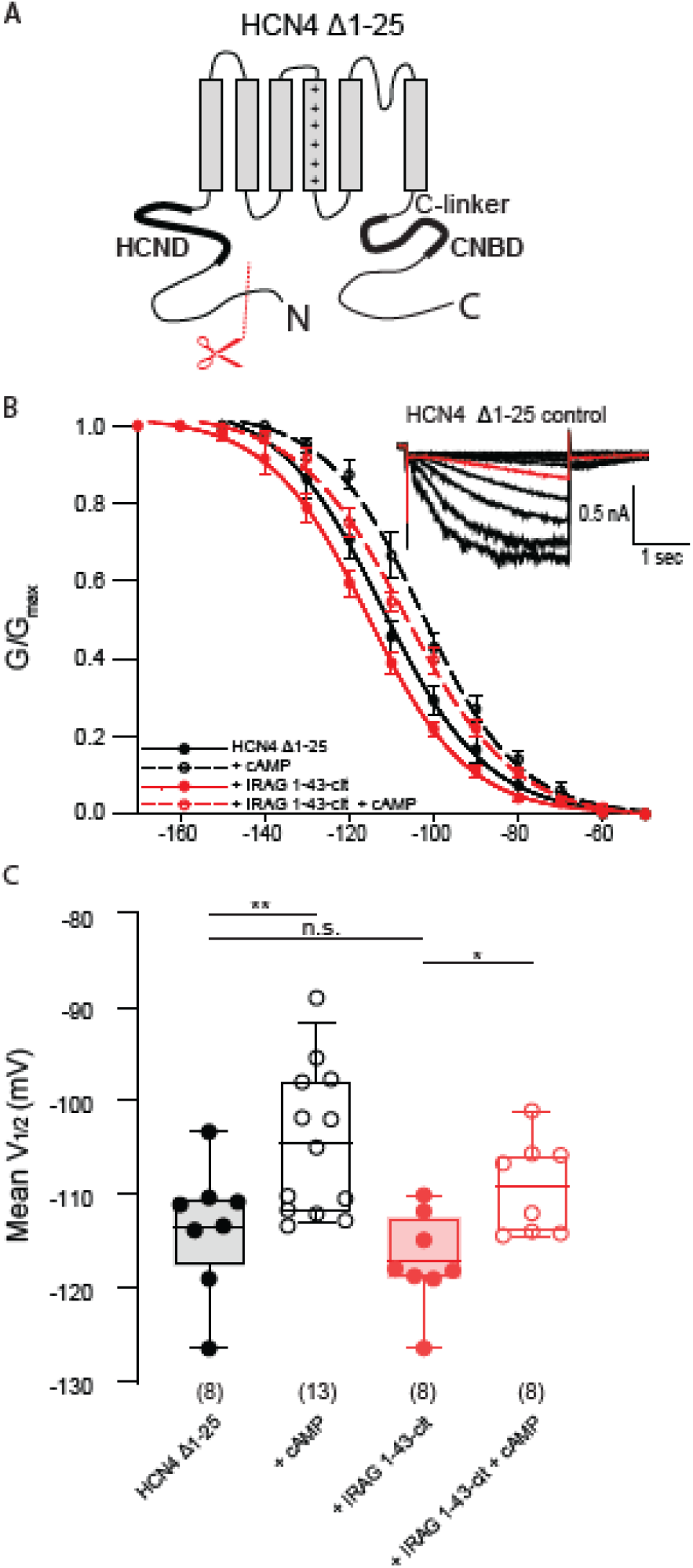
HCN4 distal N-terminus, residues 1-25, are required for regulation by IRAG 1-43. **(A)** Schematic depicting the domains of HCN4. The conserved HCND and CNBD/C-linker are represented by thicker black lines. Red scissors indicate truncation of the initial 25 residues of. **(B)** Voltage dependence of activation for HCN4 Δ1-25 in the absence (black) or presence (red) of IRAG 1-43-cit and/or 1 mM cAMP (open symbols); *inset*: Current family HCN4 Δ1-25 control. Current in red is elicited at -110 mV. Error bars represent SEM. **(C)** Box plot of mean midpoints of activation for HEK293 cells transiently expressing HCN4 Δ1-25 in the presence or absence of IRAG 1-43-cit in the absence or presence of 1 mM cAMP using the same color scheme as in B. Lines represent mean, box represents IQR, whiskers represent 10-90^th^ percentile range. * represents p < 0.05, ** represents p < 0.01, and *** represents p < 0.001. All means, standard errors, and exact p-values are in **Supplementary Table 3**.

Next, we examined the role that the unique distal C-terminus of HCN4 plays in IRAG regulation. We generated a mutant HCN4 channel missing the ∼500 amino acid C-terminal domain downstream of the CNBD (HCN4-S719X; **Fig. 4A**). Again, the truncated channel showed the normal depolarizing shift in response to 1 mM cAMP (**Fig. 4B, C**. However, co-expression with IRAG 1-43-cit resulted in only a modest rightward shift in the voltage dependence of activation. Inclusion of 1 mM cAMP in the presence of IRAG 1-43-cit produced a small but significant depolarizing shift in the V_1/2_. While certainly not conclusive, these data suggest that the non-conserved portion of the C-terminus might contribute to the ability of IRAG to modulate HCN4 gating, perhaps by stabilizing the position of the CNBD relative to the transmembrane domains of the channel.

**Figure 4.**
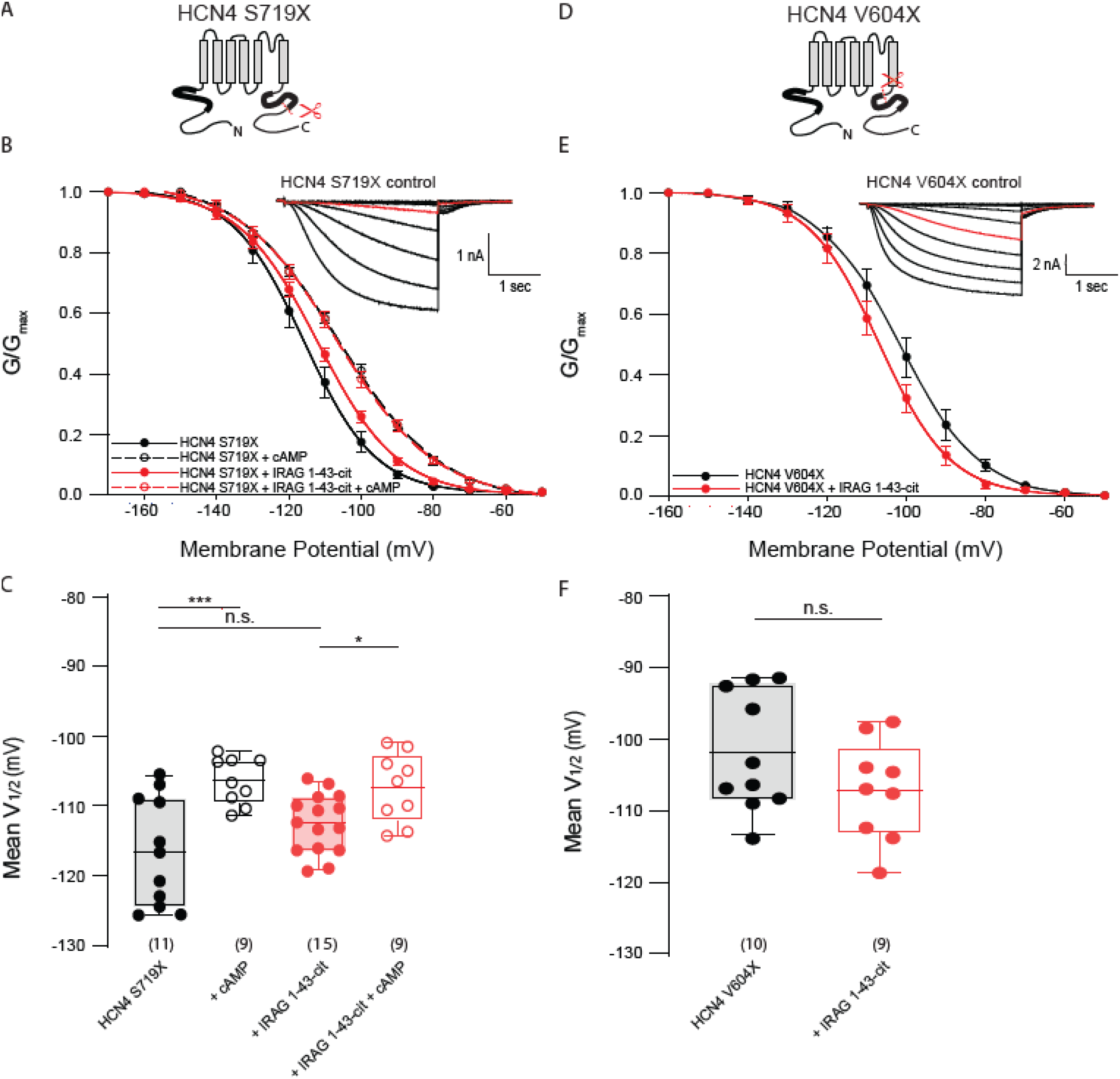
The C-terminus of HCN4 is required for regulation by IRAG 1-43. **(A)** Schematic depicting the HCN4 S719X truncation of the distal C-terminus. The conserved HCND and CNBD/C-linker are represented by thicker black lines. Red scissors indicate truncation at residue HCN4 S719. **(B)** Voltage dependence of activation for HCN4 S719X in the absence (black) or presence (red) of IRAG 1-43-cit and/or 1mM cAMP (open symbols). Error bars represent SEM. *Inset*: Current family for the control condition HCN4 S719X, current in red is elicited at -110 mV. **(C)** Box plot of mean midpoints of activation for HEK293 cells transiently expressing HCN4 S719X in the presence or absence of IRAG 1-43-cit in the absence or presence of 1 mM cAMP using the same color scheme as in B. Lines represent mean, box represents IQR, whiskers represent 10-90 percentile range. * represents p < 0.05, ** represents p < 0.01, and *** represents p < 0.001. All means, standard errors, and exact p-values are in **Supplementary Table 4**. (**D**) Schematic depicting the HCN4 V604X truncation of the distal C-terminus. The conserved HCND and CNBD/C-linker are represented by thicker black lines. Red scissors indicate truncation of the entire C-terminus at residue HCN4 V604, just after S6. **(E)** Voltage dependence of activation for HCN4-V604X in the absence (*black*) or presence (*red*) of IRAG 1-43-cit and/or 1 mM cAMP (*open symbols*). Error bars represent SEM. *Inset:* Current family for the control condition HCN4 V604X. Current in red is elicited at -110mV. **(F)** Box plot of mean midpoints of activation for HEK293 cells transiently expressing HCN4-V604X in the presence or absence of IRAG 1-43-cit in the absence or presence of 1 mM cAMP using the same color scheme as E. Lines represent mean, box represents IQR, whiskers represent 10-90 percentile range. * represents p < 0.05, ** represents p < 0.01, and *** represents p < 0.001. All means, standard errors, and exact p-values are in **Supplementary Table 5**.

We previously showed that, while IRAG shifts the voltage dependence of activation, it has no impact on the rate of channel closing and still permits cAMP to bind (Peters et al., 2020b).

These data indicate that IRAG does not function as a true cAMP mimetic, because cAMP binding slows deactivation in addition to shifting voltage dependence. However, previous studies have shown that the cAMP effects on gating and deactivation occur via separable mechanisms (Wicks et al., 2011, 2009). Thus, it is possible that IRAG acts via the CNBD to shift the V_1/2_ without affecting deactivation or cAMP binding, or that IRAG acts directly on other parts of the cyclic nucleotide transduction center, such as the HCND or the S4-S5 loop. To attempt to discriminate these possibilities, we truncated the entire C-terminus of HCN4 just downstream of S6 (HCN4-V604X; **Fig. 4D**). Since HCN4-V604X is missing the CNBD it is insensitive to cAMP. Co-expression of IRAG 1-43-cit did not produce a depolarizing shift in the voltage dependence of activation, indicating that some part(s) of the C-linker/CNBD region are required, directly or indirectly, for IRAG to modulate HCN4 (**Fig. 4E, F**).

### IRAG regulation of HCN4 requires unique residues within the cAMP transduction center

The cAMP transduction center of the family of HCN channels is largely conserved. The S4-S5 linkers are identical in the four isoforms, the HCNDs differ by only 1 residue, and the B’ helix region of the C-linker differs by only 3 residues. Although the sequences are highly conserved, the structures of HCN1 and HCN4 show subtle differences in their overall arrangement (Lee and MacKinnon, 2017; Saponaro et al., 2021) . We previously showed that mutation of two of the three unique residues in the B’ helix of HCN4 was sufficient to prevent LRMP regulation (Peters et al., 2024). To test the hypothesis that IRAG also requires these same unique residues within the cAMP transduction center of HCN4, we mutated P546 and T548 of HCN4 to their homologous residue in HCN2, A467 and F469 to produce the HCN4-PT/AF channel (**Fig. 5A, B**). We found that HCN4-PT/AF was insensitive to IRAG 1-43-cit and responded normally to cAMP in both the presence and absence of IRAG 1-43cit (**Fig. 5C, D**). This suggests that normal cyclic nucleotide modulation is intact, but IRAG requires this small unique sequence to modulate HCN4.

**Figure 5.**
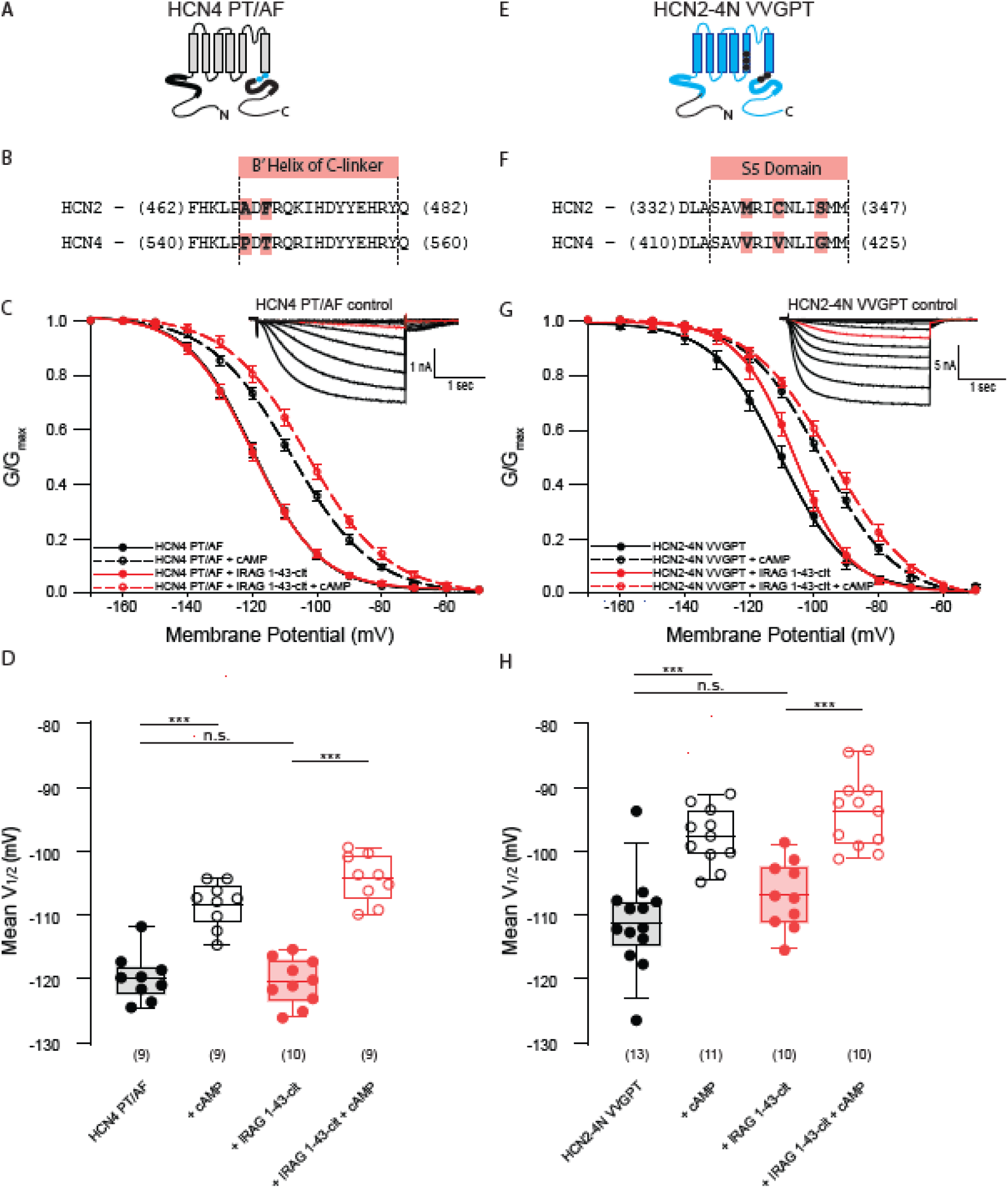
The cAMP transduction center of HCN4 is required for regulation by IRAG 1-43 and, with other unique HCN4 residues, can partially confer regulation to HCN2. **(A)** Schematic depicting the HCN4 PT/AF double mutation in the C-linker. Black background and dots represent HCN4 sequence; blue represents HCN2. **(B)** Sequence alignment of the B’ helices of the C-linker of HCN2 and HCN4. The two non-conserved residues are highlighted in light red. **(C)** Voltage dependence of activation for HCN4 PT/AF in the absence (*black*) or presence (*red*) of IRAG 1-43-cit and/or 1 mM cAMP (*open symbols*). Error bars represent SEM. *Inset:* Current family for the control condition HCN4 PT/AF. Current in red is elicited at -110 mV. **(D)** Box plot of mean midpoints of activation for HEK293 cells transiently expressing HCN4 PT/AF in the presence or absence of IRAG 1-43-cit in the absence or presence of 1 mM cAMP using the same color scheme as C. Lines represent mean, box represents IQR, whiskers represent 10-90 percentile range. * represents p < 0.05, ** represents p < 0.01, and *** represents p < 0.001. All means, standard errors, and exact p-values are in **Supplementary Table 6. (E)** Schematic depicting the HCN2-4N VVGPT chimera. Black background and dots represent HCN4 sequence; blue represents HCN2. **(F)** Sequence alignment of the S5 transmembrane helices of HCN2 and HCN4. The three non-conserved residues are highlighted in red. **(G)** Voltage dependence of activation for HCN2-4N VVGPT in the absence (*black*) or presence (*red*) of IRAG 1-43-cit and/or 1 mM cAMP (*open symbols*). Error bars represent SEM. *Inset:* Current family for the control condition HCN2-4N VVGPT. Current in red is elicited at -110 mV. **(H)** Box plot of mean midpoints of activation for HEK293 cells transiently expressing HCN2-4N VVGPT in the presence or absence of IRAG 1-43-cit and in the absence or presence of 1 mM cAMP using the same color scheme as G. Lines represent mean, box represents IQR, whiskers represent 10-90 percentile range. * represents p < 0.05, ** represents p < 0.01, and *** represents p < 0.001. All means, standard errors, and exact p-values are in **Supplementary Table 7**.

We were previously able to confer LRMP regulation to HCN2 channels in a multidomain chimera (HCN2-4N-VVGPT) in which the N-terminus of HCN2 was replaced by the HCN4 N-terminus and the unique C-linker residues and the three unique S5 residues were mutated to the cognate HCN4 residues (Peters et al., 2024) (**Fig. 5E, F**). HCN2-4N-VVGPT responded to cAMP with an ∼13.4mV depolarizing shift in the voltage dependence of activation (**Fig. 5G, H**). Co-expression of IRAG 1-43-cit with HCN2-4N VVGPT produced only a modest rightward shift in the voltage-dependence of activation, suggesting that while this region is critical for the IRAG effect, these mutations are not sufficient to confer the effect to HCN2. Addition of 1 mM cAMP to cells expressing both the mutant channel and IRAG 1-43-cit produced a significant shift in the V_1/2_. These data are consistent with limited regulation of HCN2-4N VVGPT by IRAG 1-43.

## DISCUSSION

In this study, we show that IRAG regulates HCN4 via the first 43 residues of the IRAG N-terminus as well as multiple domains of the channel. These findings are consistent with a model in which IRAG exerts its gain-of-function effect on HCN4 via isoform-specific features of the intramolecular cAMP transduction pathway of the channel (Saponaro et al., 2021; Porro et al., 2019; Wang et al., 2020; Weißgraeber et al., 2017; Kondapuram et al., 2022). These findings are analogous to, though opposite in polarity from, the mechanism of action of LRMP on HCN4. Given previous studies showing that IRAG scaffolds signaling proteins and that it is highly expressed in the sinoatrial node of the heart, the work has implications for the regulation of cardiac pacemaking.

### IRAG and LRMP

Our present finding that the initial N-terminal 43 amino acids of IRAG are necessary and sufficient for regulation of HCN4 echoes our previous discovery that the distal N-terminus of LRMP is also required for regulation of HCN4 (Peters et al., 2024). However, this similarity may represent more of a coincidence of convenience rather than a true commonality of mechanism: the N-terminal domains of LRMP and IRAG can in principle extend the farthest toward the plasma membrane from their C-terminal ER anchors, making them logical candidates for interaction with channels on the plasma membrane despite the fact that they exhibit significant sequence divergence. Alignment of the initial 43 residues of IRAG with the initial 227 residues of LRMP does not reveal any obvious common motifs that would suggest a conserved binding site.

In our previous study of LRMP regulation of HCN4, we also demonstrated physical association between the N-terminals of LRMP and HCN4 via Förster Resonance Energy Transfer (FRET). Although we attempted similar FRET experiments with many different fragments of IRAG and HCN4, none of the constructs tested — including residues 1-43 of IRAG — exhibited FRET above baseline with any domain of HCN4. This could be due to a relatively low affinity interaction between IRAG and fragments of HCN4 that the FRET cannot measure. It might require the full three dimensional structure of the channel to create a higher affinity site for IRAG binding. However, the regulation of HCN4 by residues 1-43 of IRAG is robust and does not differ from regulation by the full-length IRAG, whether the fragment is expressed as a transfected plasmid or introduced as a peptide in the patch pipette. These observations are consistent with a relatively fast, direct interaction of IRAG1-43 with the channel, as we saw for LRMP, rather than an indirect action involving transcriptional or translational mechanisms.

On the channel side of the regulation by IRAG, we found that the initial 25 residues of the N-terminus of HCN4 are required. We chose to truncate HCN4 at residue 25 because this location is an alternative translation initiation site. HCN4 channels lacking residues 1-25 were previously reported to be insensitive to cAMP in excised inside-out patches (Liu and Aldrich, 2011), however our data indicate a significant cAMP response in HCN4Δ1-25 in whole cell recordings. We suspect that the discrepancy in results arises from differences in recording conditions, such as free Ca^2+^ levels in solutions, reflecting the sensitivity of this area to regulation, but more work would be needed to examine these differences in detail. The conserved proximal N-terminal domain of HCN channels forms the so-called HCN domain, which is positioned close to the C-linker domain and S4-S5 linker in the structure (Lee and MacKinnon, 2017). It is thus possible that the unique distal N-terminus of HCN4 contributes to regulation by IRAG by altering relationships within this “transduction center.”

In addition to the distal N-terminus, we also found that the C-terminus and unique regions within the channel’s putative intramolecular cAMP transduction pathway also contribute to the regulation by IRAG. Since the chimera HCN2-4N VVGPT conferred to HCN2 sensitivity to LRMP but not IRAG, it is possible that additional components of HCN4 may be necessary for IRAG regulation as compared to LRMP. However, it’s not clear that additional chimeras would be meaningful given the involvement of multiple domains and the high conservation of residues within the intracellular regions of the channels. Meanwhile, the involvement of multiple sites along the pathway that links the CNBD to channel gating suggests that IRAG, like LRMP, may act by modifying the strength of coupling between the CNBD and gate of HCN4. The unliganded CNBD of HCN channels exerts a negative effect on gating, known as autoinhibition. Binding of cAMP relieves the autoinhibition to cause the canonical depolarizing shift in the voltage-dependence of activation. Our data thus suggest that LRMP could act to stabilize the unliganded state of the CNBD and intramolecular transduction pathway, whereas IRAG could exert its depolarizing influence by stabilizing a state more similar to the liganded state. Ultimately, understanding the mechanisms of action of LRMP and IRAG will benefit from structures of complexes with HCN4. At present, no structural information is available for either LRMP or IRAG. Moreover, protein structure prediction algorithms produce only low confidence predictions for largely unstructured regions in the N-terminal domains of LRMP and IRAG which preclude even rudimentary docking attempts.

### Potential physiological significance in the sinoatrial node (SAN)

Cardiac pacemaking in the sinoatrial node of the heart involves spontaneous changes in membrane potential in specialized pacemaker cells. These changes in membrane potential are driven directly by the gating of voltage-gated ion channels, including HCN4, and indirectly by intracellular Ca^2+^ signaling, which alters membrane potential primarily via activity of the electrogenic Na-Ca^2+^ exchanger. Although both the so-called “membrane clock” and “Ca^2+^ clock” are critical for pacemaking, no molecular mechanisms have yet been identified to coordinate the activity of these two systems. HCN4 is a molecular marker of the sinoatrial node and is critical for cardiac pacemaking (Moosmang et al., 2001; Stieber et al., 2003; Schulze-Bahr et al., 2003; Baruscotti et al., 2011; Peters et al., 2021). IRAG is an ER/SR transmembrane protein that is expressed in the SAN at levels similar to that of HCN4 (Peters et al., 2020a). In other cell types, IRAG is known to regulate ER/SR Ca^2+^ release via the IP3 receptor. Hence, the interaction between HCN4 and IRAG could serve as a molecular bridge between changes in membrane potential and intracellular Ca^2+^ release.

IRAG is also well-established component of the IP_3_R-cGMP signaling pathway, where it acts as a molecular scaffold to organize proteins involved in cGMP signaling(Schlossmann et al., 2000; Ammendola et al., 2001). The involvement of the N-terminal domains of LRMP and IRAG in regulation of HCN4 allows the possibility that the coiled-coil domains of the two proteins may orient signaling molecules, such as cGMP-dependent protein kinase (PKG), in close proximity to the channel’s gating machinery. Upon increases in cGMP, tethered PKG1beta phosphorylates IRAG at multiple sites, which could induce conformational changes in the N-terminal region and its interaction with HCN4(Geiselhöringer et al., 2004b). It is thus possible that PKG signaling could provide additional nuance or range to the regulation of HCN4 by IRAG.

### Conclusions

In conclusion, we have shown that isoform-specific regulation of HCN4 by IRAG requires the first 43 residues of IRAG and multiple domains of HCN4 along the intramolecular cAMP transduction pathway. Although IRAG and its homolog LRMP exert opposite effects on HCN4, they appear to work via similar intramolecular pathways, perhaps by stabilizing/destabilizing the inhibitory effects of the unliganded CNBD. The high expression of IRAG in the sinoatrial node of the heart and its known interactions with cGMP signaling molecules suggests that it could contribute to regulation of cardiac pacemaking.

## DATA AVAILABILITY STATEMENT

The data files supporting all findings presented in this paper are available from the corresponding author.

## ACKNOWLEDGEMENTS

Research reported in this publication was supported by the National Institute of General Medical Sciences R35GM137912 (to JRB), R01GM140004 (to CP), and R01HL088427 (to CP). The content is solely the responsibility of the authors and does not necessarily represent the official views of the National Institutes of Health or National Science Foundation.

## Author contributions

JB and CP: conceptualization, data curation, methodology, project administration, supervision, resources, and writing—original draft, review, and editing. LMB: data curation, methodology, formal analysis, writing—original draft, review, and editing. CHP, ET, RR, and WGN: conceptualization, data curation, methodology. DJE and AAL: resources and methodology.

## ABBREVIATIONS

cAMP: cyclic Adenosine Monophosphate
LRMP: Lymphoid Restricted Membrane Protein
IRAG: Inositol triphosphate Receptor Associated Guanylate kinase substrate
FRET: Forster Resonance Energy Transfer
HCN: Hyperpolarization activated, Cyclic-Nucleotide sensitive channels
CNBD: Cyclic Nucleotide Binding Domain
HCND: HCN Domain

## SUPPLEMENTARY TABLES

**Supplementary Table 1:** Two-way ANOVA with Bonferroni post hoc test for multiple comparisons. Mean values are reported ± SEM with n listed in parentheses. P values below columns represent the comparison between the two above values; P values at the end of rows represent the comparison between the two values to the left.

| 2-Way ANOVA |  |  |  |
| --- | --- | --- | --- |
| condition | Control | IRAG | IRAG vs. Control |
| HCN4 | -117.1 ± 1.4 (15) | -104.3 ± 2.3 (11) | p < 0.001 |
| + cAMP | -104.8 ± 1.4 (8) | -105.0 ± 2.8 (10) | p = 0.954 |
|  | p < 0.001 | p = 0.829 |  |

**Supplementary Table 2:** Two-way ANOVA with Bonferroni post hoc test for multiple comparisons. Mean values are reported ± SEM with n listed in parentheses. P values below columns represent the comparison between the two above values; P values at the end of rows represent the comparison between the two values to the left.

| 2-Way ANOVA |  |  |  |  |  |  |  |
| --- | --- | --- | --- | --- | --- | --- | --- |
| condition | control | IRAG Δ1-43 | vs. control | IRAG 1-43-cit | vs. control | 10uM IRAG 1-43 | vs. control |
| HCN4 WT | -117.1 ± 1.4 (15) | -116.1 ± 0.9 (9) | p = 1.000 | -109.2 ± 2.3 (16) | p = 0.008 | -103.9 ± 4.4 (7) | p < 0.001 |
| + cAMP | -104.8 ± 1.4 (8) | -101.9 ± 1.5 (9) | p = 1.000 | -105.1 ± 1.2 (11) | p = 1.000 | -105.8 ± 2.7 (7) | p = 1.000 |
|  | p < 0.001 | p < 0.001 |  | p = 0.130 |  | p = 0.598 |  |

**Supplementary Table 3:** Two-way ANOVA with Bonferroni post hoc test for multiple comparisons. Mean values are reported ± SEM with n listed in parentheses. P values below columns represent the comparison between the two above values; P values at the end of rows represent the comparison between the two values to the left.

| 2-Way ANOVA |  |  |  |
| --- | --- | --- | --- |
| condition | control | IRAG 1-43-cit | IRAG vs. Control |
| HCN4 Δ1-25 | -113.6 ± 2.4 (8) | -117.2 ± 1.8 (8) | p = 0.281 |
| + cAMP | -104.6 ± 2.2 (13) | -109.2 ± 1.8 (8) | p = 0.127 |
|  | p = 0.005 | p = 0.022 |  |

**Supplementary Table 4:** Two-way ANOVA with Bonferroni post hoc test for multiple comparisons. Mean values are reported ± SEM with n listed in parentheses. P values below columns represent the comparison between the two above values; P values at the end of rows represent the comparison between the two values to the left.

| 2-Way ANOVA |  |  |  |
| --- | --- | --- | --- |
| condition | control | IRAG 1-43-cit | IRAG vs. Control |
| HCN4 S719X | -116.6 ± 2.4 (11) | -112.5 ± 1.1 (15) | p = 0.062 |
| + cAMP | -106.5 ± 1.1 (9) | -107.4 ± 1.7 (9) | p = 0.709 |
|  | p < 0.001 | p = 0.030 |  |

**Supplementary Table 5:** Two-tailed t-test between the two conditions. Mean values are reported ± SEM with n listed in parentheses. P value is reported in the final column.

| two-tailed t-test |  |  |  |
| --- | --- | --- | --- |
| condition | control | IRAG 1-43-cit | IRAG vs. Control |
| HCN4 V604X | -101.9 ± 2.6 (10) | -107.1 ± 2.3 (9) | p = 0.160 |

**Supplementary Table 6:** Two-way ANOVA with Bonferroni post hoc test for multiple comparisons. Mean values are reported ± SEM with n listed in parentheses. P values below columns represent the comparison between the two above values; P values at the end of rows represent the comparison between the two values to the left.

| 2-Way ANOVA |  |  |  |
| --- | --- | --- | --- |
| condition | control | IRAG 1-43-cit | IRAG vs. Control |
| HCN4 PT/AF | -119.8 ± 1.3 (9) | -120.5 ± 1.2 (10) | p = 0.671 |
| + cAMP | -108.3 ± 1.2 (9) | -104.3 ± 1.3 (9) | p = 0.026 |
|  | p < 0.001 | p < 0.001 |  |

**Supplementary Table 7:**
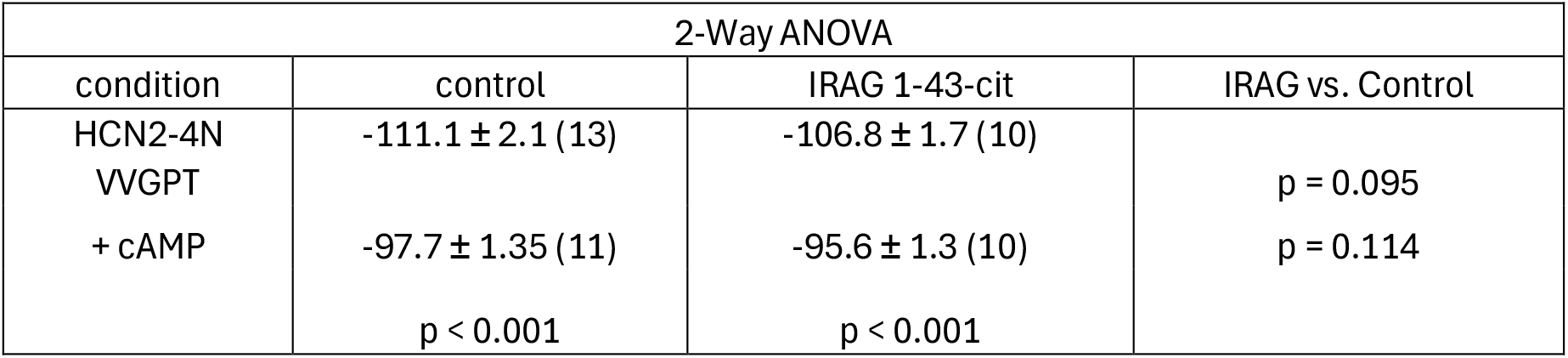
Two-way ANOVA with Bonferroni post hoc test for multiple comparisons. Mean values are reported ± SEM with n listed in parentheses. P values below columns represent the comparison between the two above values; P values at the end of rows represent the comparison between the two values to the left.

